# Mathematical Modeling of Estrogen-Mediated Inflammation in the Gut and Immune System

**DOI:** 10.1101/2024.12.23.630175

**Authors:** Ariel M. Lighty, Mohammad Aminul Islam, Brenda J. Smith, Ashlee N. Ford Versypt

**Affiliations:** Department of Chemical and Biological Engineering, University at Buffalo, The State University of New York, Buffalo, New York, United States of America; Indiana Center for Musculoskeletal Health, Indiana School of Medicine, Indianapolis, IN 46202, USA; Department of Obstetrics and Gynecology, Indiana School of Medicine, Indianapolis, IN 46202, USA; Department of Biomedical Engineering, University at Buffalo, The State University of New York, Buffalo, New York, United States of America; Institute for Artificial Intelligence and Data Science, University at Buffalo, The State University of New York, Buffalo, New York, United States of America; Department of Pharmaceutical Sciences, University at Buffalo, The State University of New York, Buffalo, New York, United States of America

**Keywords:** mathematical modeling, inflammation, sex differences, T cells, immunology, gut immunity, aging, cell differentiation, osteoporosis

## Abstract

Postmenopausal osteoporosis is a chronic inflammatory disease characterized by decreased bone mass and increased bone fracture risk. Estrogen deficiency during menopause plays a major role in post-menopausal osteoporosis by influencing bone, immune, and gut cell activity. In the gut, estrogen loss decreases tight junction proteins that bind epithelial cells of the intestinal barrier together. This increases gut permeability and alters the balance of T cells, increasing inflammatory T helper (Th17) cells and decreasing anti-inflammatory regulatory T cells (Tregs). Prebiotic dietary fibers are a promising treatment modality for gut-mediated inflammation and bone loss associated with estrogen deficiency. In a previous study, we investigated the effect of butyrate (a prebiotic by-product) on Treg levels in the gut, blood, and bone starting with healthy conditions. However, a gap remains in our mechanistic understanding of how estrogen affects systemic inflammation separately and in combination with prebiotic fibers.

We developed a two-compartment dynamic model of estrogen in the gut and immune system to investigate the mechanisms of gut-mediated inflammation induced by estrogen-deficient conditions. Our initial model contains ODEs for estrogen, naive T cells, Tregs, and Th17 cells in gut and blood compartments. Estrogen is modeled using physiological mass balance equations considering its production, migration, and elimination. We integrate our estrogen model with an adapted version of our previous butyrate model, adding a population balance for Th17 cells and terms for bidirectional migration of Tregs and Th17 cells. Estrogen interactions with cells are modeled using Hill-type activation and inhibition functions. Our mathematical model predicts changes in T cell dynamics associated with changes in endogenous estrogen levels. We compare our predictions to in vivo results of estrogen, Tregs, and Th17 levels in the gut and blood after surgical menopause in mice. In the future, we will leverage this model to investigate whether the effect of prebiotic fiber treatment on estrogen-deficient bone loss is significantly affected by gut-mediated inflammation. The emphasis in this preprint is on the methods, particularly those for literature-based model parameterization.

## 1 Introduction

Postmenopausal osteoporosis (PMO) is a chronic inflammatory disease characterized by decreased bone mass and increased bone fracture risk. Estrogen deficiency during menopause plays a major role in PMO by influencing bone, immune, and gut cell activity [1, 2]. In the gut, estrogen loss decreases tight junction proteins that bind epithelial cells of the intestinal barrier together [3]. This increases gut permeability and alters the balance of T cells, increasing inflammatory T helper (Th17) cells and decreasing anti-inflammatory regulatory T cells (Tregs) [3]. Prebiotic dietary fibers are a promising treatment modality for gut-mediated inflammation and bone loss associated with estrogen deficiency [4, 5]. In a previous study, we investigated the effect of butyrate (a prebiotic by-product) on Treg levels in the gut, blood, and bone starting with healthy conditions [6]. However, a gap remains in our mechanistic understanding of how estrogen affects systemic inflammation separately and in combination with prebiotic fibers.

We developed a two-compartment model of estrogen in the gut and blood system to investigate the mechanisms of gut-mediated inflammation induced by estrogen-deficient conditions on T cell dynamics (Figure 1). Our initial model contains ODEs for estrogen, naïve T cells, Tregs, and Th17 cells in gut and blood compartments. Estrogen is modeled using physiological mass balance equations that consider its production, migration, and elimination. We integrate our estrogen model with an adapted version of our previous butyrate model [6], where we add a population balance for Th17 cells and terms for bidirectional migration of Tregs and Th17 cells. Estrogen interactions with cells are modeled using Hill-type activation and inhibition functions. Our mathematical model predicts changes in T cell dynamics associated with changes in endogenous estrogen levels. We compare our predictions to in vivo results of estrogen, Tregs, and Th17 levels in the gut and blood after surgical menopause in mice [7, 8]. In the future, we will leverage this model to investigate whether the effect of prebiotic fiber treatment on estrogen-deficient bone loss is significantly affected by gut-mediated inflammation.

**Figure 1:**
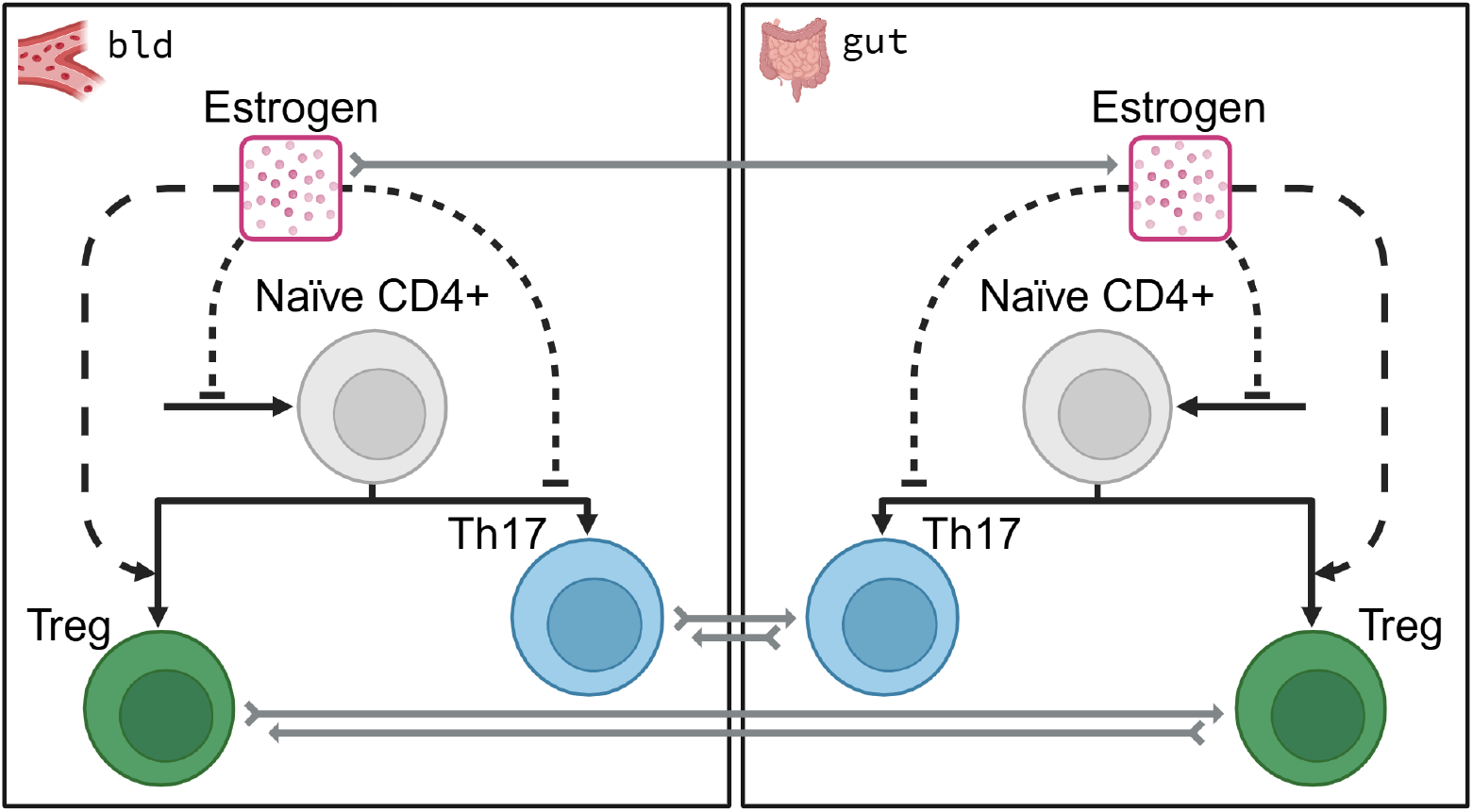
A two-compartment model of estrogen effects on T cells in the blood and gut. Estrogen migrates from the blood (bld) compartment to the gut (gut) compartment. Tregs and Th17 cells migrate between the gut and blood compartments. In both compartments, estrogen inhibits the formation of naïve CD4^+^T cells, stimulates naïve CD4^+^differentiation to Tregs, and inhibits naïve CD4^+^differentiation to Th17 cells. Cell populations tracked are illustrated as circles with a nucleus (Treg in green, Th17 in blue, and naïve CD4^+^in gray). Estrogen, the only chemical species tracked, is illustrated by pink dots in a square. Solid black arrows denote differentiation of one cell to another, and solid gray arrows denote cell migration from one compartment to another. Dotted black lines/arrows denote interactions that influence processes without species production or consumption, where arrowhead endings denote process stimulation (up-regulation) and flathead endings denote process inhibition (down-regulation).

## 2 Methods

### 2.1 Estrogen Model Formulation

The following section describes model formulations in which estrogen dynamics are decoupled from T cell dynamics. The models are decoupled so that we can explore how a constant or exponential decay in estrogen affects T cell dynamics independently.

Estradiol is the dominant estrogen during reproductive years [9], so it is a reasonable measure for understanding the effects of estrogen before and after menopause or OVX. E in the equations defined below thus represents estradiol levels, rather than total estrogen levels.

We formulate a physiologically-based model from the following assumptions:

1. Estrogen levels in the gut are independent of ovarian secretion [7]
2. Estrogen ‘production’ in the blood is dependent on ovarian secretion rates
3. Estrogen migrates primarily from the blood into the gut
4. Estrogen migration from the gut into the blood is negligible (only 5% of estrogen produced in non-ovarian tissue enters circulation [9])
5. The removal/excretion of estrogen can be capture by its half-life

The resulting equations are:

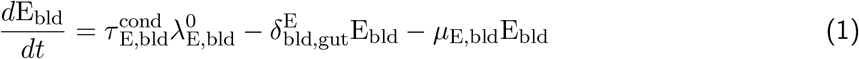

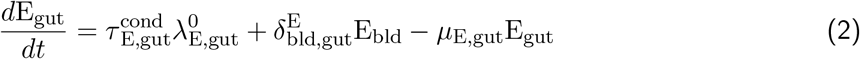

where 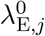 is the influx or production of estrogen in compartment *j* and 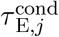 modifies estrogen influx or production rates based on the state of ovarian function in compartment *j* under sham or OVX surgery conditions cond, while 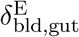 is the migration of estrogen from the blood into the gut and *µ*_E,j_ is the half-life of estrogen in compartment *j*.

#### 2.1.1 Parameters and initial conditions

The initial conditions E_bld,0_ and E_gut,0_ are set equal to the estrogen levels of sham surgery mice in Oakley et al. [7] (see Section 2.8.9). Initial conditions and parameter values are outlined in Table 1.

**Table 1:** Estrogen model initial conditions and parameters.

| Symbol | Value | Unit | Source |
| --- | --- | --- | --- |
| $E_{\text{bld},0}$ | $0.151 \pm 0.014$ | pg | [7] |
| $E_{\text{gut},0}$ | $14.79 \pm 4.58$ | pg | [7] |
| $\tau_{E,\text{bld}}^{\text{sham}}$ | 1 | – | Assumption |
| $\tau_{E,\text{gut}}^{\text{sham}}$ | 1 | – | Assumption |
| $\lambda_{E,\text{bld}}^0$ | $16.6 \pm 2.7$ | pg/day | [10] |
| $\lambda_{E,\text{gut}}^0$ | $0.64 \pm 0.12$ | pg/day | [7] |
| $\mu_{E,\text{bld}}$ | 4.4 | day<sup>-1</sup> | Bauss1996 |
| $\delta_{\text{bld,gut}}^E$ | 105 | day<sup>-1</sup> | Fitted |
| $\mu_{E,\text{gut}}$ | 1.12 | day<sup>-1</sup> | Fitted |
| $\tau_{E,\text{bld}}^{\text{ovx}}$ | 0.55 | – | Fitted |
| $\tau_{E,\text{gut}}^{\text{ovx}}$ | 8.8 | – | Fitted |

Estrogen model B contains 2 equations with 7 parameters: 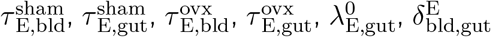 and *µ*_E_. Estrogen half-life in the blood *µ*_E,bld_ and production rates in both compartments 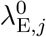 are estimated from the literature.

The values for estrogen half-life in the gut *µ*_E,gut_ and migration into the gut 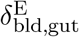 are calibrated to fit the healthy condition. We assume the healthy or sham estrogen levels are the steady state levels.

That is, we assume 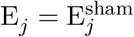 when *d*E_j_*/dt* = 0.

The 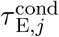 parameters are defined by the ovarian function. We assume healthy ovaries are represented by 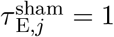. Reduced ovarian function, specifically ovary removal, is represented by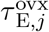. The values for 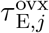 are calibrated to fit OVX estrogen levels.

### 2.2 T Cell Model Formulation

#### 2.2.1 Adapting a model of butyrate effects on T cells

General assumptions for constructing Figure 1:

1. There are no changes in butyrate from homeostasis
2. Estrogen inhibits the formation of naïve T cells
3. Estrogen inhibits naïve T cell differentiation to Th17 cells
4. Estrogen stimulates naïve T cell differentiation to Treg cells

We begin developing our model of T cell dynamics based on Islam et al. [6]. This work develops a 3-compartment model that tracks the dynamics of butyrate, CD4^+^naïve T cells, and Tregs in the gut, blood, and bone. Equations for the gut and blood compartments are shown below.

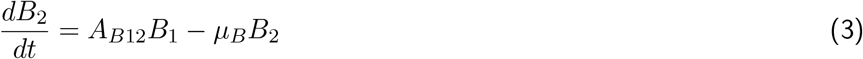

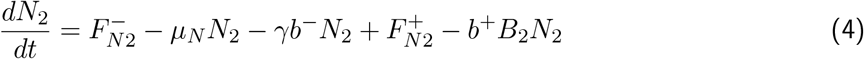

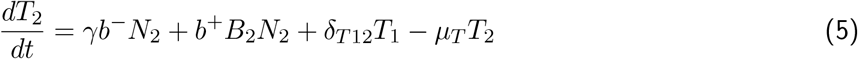

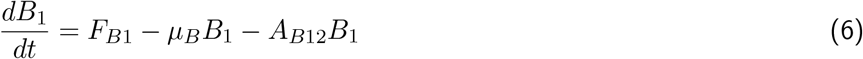

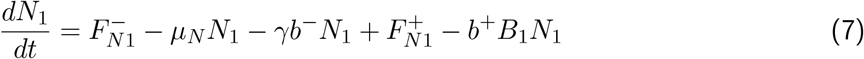

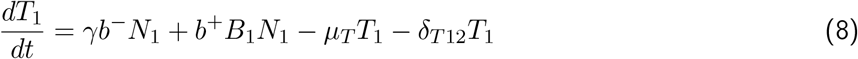

Equations 6–8 *track changes in butyrate (B*) concentration, naïve T cell (*N*) populations, and Treg populations (*T*) relative to homeostasis in the gut and blood. The circulation of naïve T cells is not considered. It was also assumed removal rates (*µ*_*i*_) and naïve T cell homeostatic differentiation rates 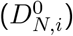 were equivalent in the gut and blood compartment.

The model tracks changes in butyrate as a deviation variable. That is, it tracks changes in butyrate from the homeostatic condition, which is defined as no butyrate supplement. So the initial condition of butyrate is zero. This is represented mathematically as *B*_1_ = *B*_2_ = *B*_3_ = 0 at *t* = 0. At steady state, the value of *B*_*j*,ss_ is the amount of excess butyrate in compartment *j* due to treatment.

Since T cell populations are typically reported as a percent of total naïve CD4^+^T cells in experimental literature, an arbitrary basis of *N*_*j*_(*t* = 0) = *N*_*j*,0_ = 1 is chosen for initial homeostatic naïve T cell counts. This allows us to assume the initial fraction of Tregs 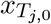 per compartment *j* is given by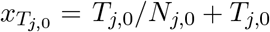. Rearranging,

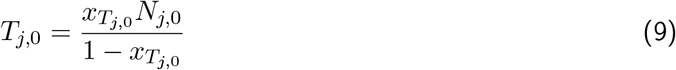

gives an equation for the initial number of Tregs.

The parameters are defined as follows:

▪ *F*_*Bj*_ = zero-order source term for butyrate
▪ *µ*_*B*_ = lumped degradation rate term that includes degradation, local tissue loss, and excretion
▪ *A*_*Bji*_ = absorption rate from compartment *j* into compartment *i*
▪ 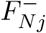= formation rate of naïve T cells in compartment *j* under homeostatic conditions, i.e., no excess butyrate
▪ *γb*^−^ = differentiation rate of naïve T cells under homeostatic conditions to Treg cells
▪ *γ* = coefficient for similarity to Treg polarizing conditions
▪ *b*^−^ = homeostasis differentiation rate constant of naïve T cells to Tregs
▪ *µ*_*N*_ = degradation rate for naïve T cells
▪ 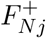= increase in formation rate of naïve T cells in compartment *j* due to stimulation by excess butyrate
▪ *b*^+^ = increase in differentiation rate of naïve T cells to Treg cells due to stimulation by excess butyrate
▪ *γb*^−^ = differentiation rate of naïve T cells under homeostatic conditions to Treg cells
▪ *b*^+^ = increase in differentiation rate of naïve T cells to Treg cells due to stimulation by excess butyrate
▪ *µ*_*T*_ = degradation rate of Treg cells
▪ *δ*_*T ji*_ = migration rate of Treg cells from compartment *j* to compartment *i*

#### 2.2.2 Re-configuring variable names

Estradiol or 17β-estradiol is often abbreviated as E2. For added clarity, we therefore relabel the compartments from 1 to gut and 2 to bld. Next, we denote Tregs as R rather than *T* since we are interested in adding Th17 (to be denoted H) dynamics. The following variable modifications are also made:

▪ *A*_*Bji*_ → 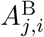
▪ 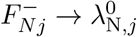
▪ *γ* → *γ*^R^
▪ *b*^−^ → 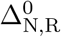
▪ 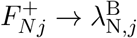
▪ *b*^+^ → 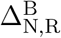
▪ *δ*_*T ji*_ →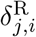

where *j* and *i* are compartments. Equations 6–9 are rewritten with these new variables in Equations 10– 16.

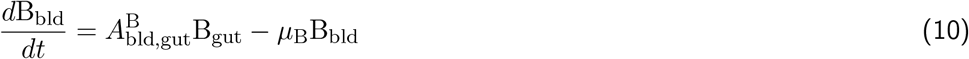

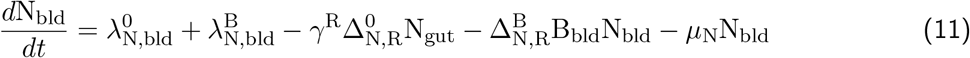

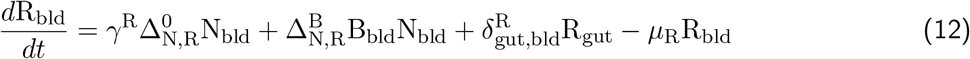

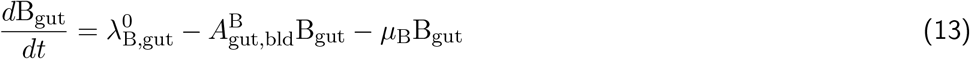

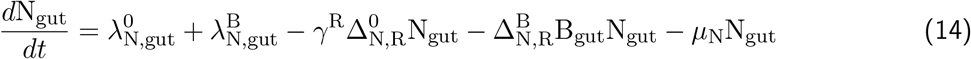

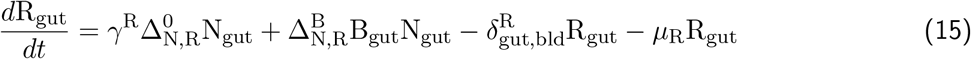

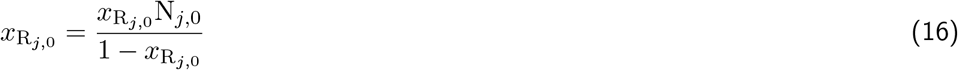

#### 2.2.3 Simplifying T cell dynamics

To adapt the model for estrogen effects, we assume no change in butyrate with changes in estrogen such that *dB*_*j*_/*dt* = 0. As a result, Equations 13–12 become

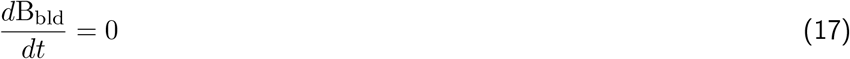

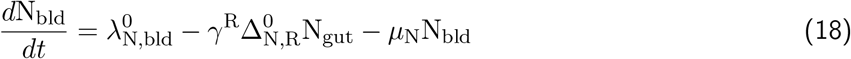

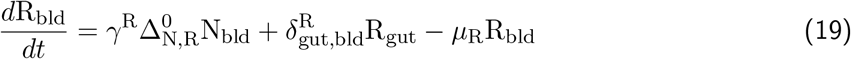

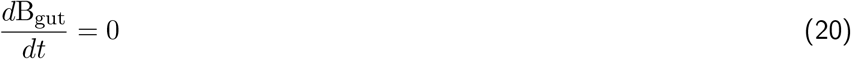

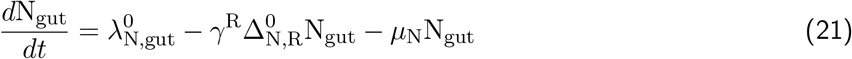

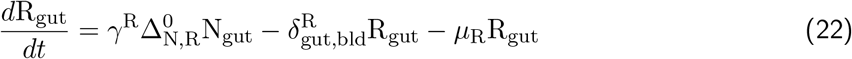

which is a simplified population model of naïve T cells and Tregs without butyrate effects.

The parameter *γ*^R^ was included in the butyrate model to account for the difference between *in vivo* and *in vitro* differentiation conditions because 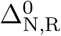 had been fit to *in vitro* data. In this model, we let *γ*^R^ = 1 by fitting 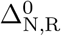 to *in vivo* data instead. The resulting T cell equations are given by Equations 25–24.

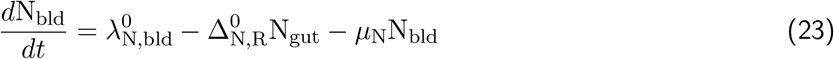

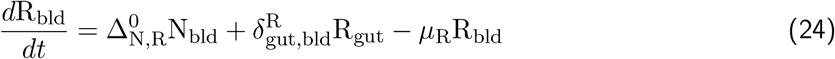

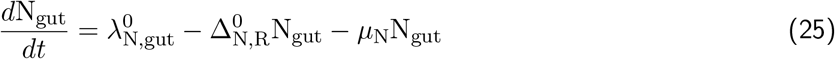

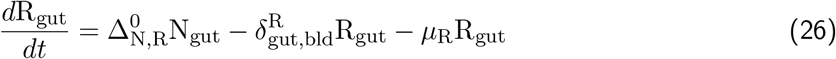

#### 2.2.4 T cell model parameters

The current T cell model formulation (Equations 36–41) contains 6 equations with 6 unknown parameters: 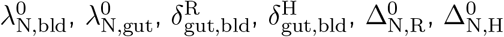.

The ratios *K*_N_ and 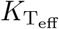 will be calculated from literature values.

The parameters *µ*_N_, *µ*_R_, and *µ*_H_ will be estimated based on the literature.

The parameters 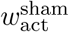 and 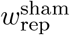 are assumed to be 1. 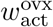 and 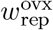 are fitted to H_bld_ and R_gut_ levels during OVX.

T cell model parameters are summarized in Table 2 below.

**Table 2:** T cell model parameters.

| Symbol | Value | Unit | Source |
| --- | --- | --- | --- |
| $w_{\text{act}}^{\text{sham}}$ | 1 | – | Assumed |
| $w_{\text{rep}}^{\text{sham}}$ | 1 | – | Assumed |
| $K_{T_{\text{eff}}}$ | 31 | – | [11] |
| $K_N$ | $0.76 \pm 0.12$ | – | [12] |
| $\mu_N$ | $2.6 \times 10^{-3}$ | day<sup>-1</sup> | [13] |
| $\mu_R$ | $4.6 \times 10^{-3}$ | day<sup>-1</sup> | [14] |
| $\mu_H$ | $4.6 \times 10^{-3}$ | day<sup>-1</sup> | [15] |
| $\lambda_{N,\text{bld}}^0$ | $6.09 \times 10^{-2}$ | cells/day | Fitted |
| $\lambda_{N,\text{gut}}^0$ | $4.62 \times 10^{-2}$ | cells/day | Fitted |
| $\Delta_{N,R}^0$ | $6.6 \times 10^{-3}$ | day<sup>-1</sup> | Fitted |
| $\Delta_{N,H}^0$ | $2.3 \times 10^{-3}$ | day<sup>-1</sup> | Fitted |
| $\delta_{\text{gut,bld}}^R$ | $2.74 \times 10^{-4}$ | day<sup>-1</sup> | Fitted |
| $\delta_{\text{gut,bld}}^H$ | $4.2 \times 10^{-3}$ | day<sup>-1</sup> | Fitted |
| $w_{\text{act}}^{\text{ovx}}$ | 0.56 | – | Fitted |
| $w_{\text{rep}}^{\text{ovx}}$ | 1.39 | – | Fitted |

**Table 3:**
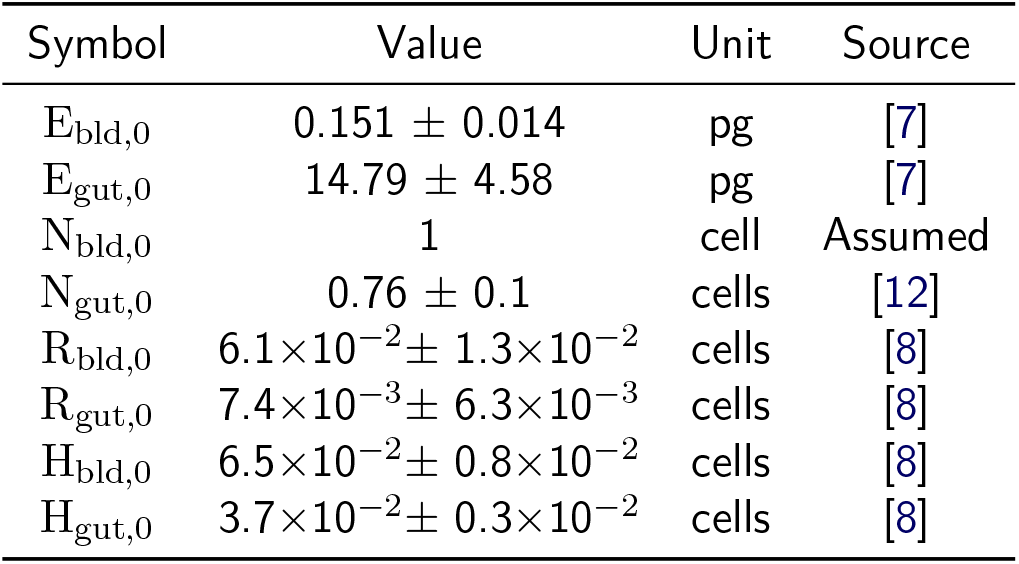
Estrogen and T cell model initial conditions.

### 2.3 Estrogen effects and Th17 cell dynamics

Estrogen (E) inhibits naïve T cells formation and their differentiation to inflammatory Th17 cells [3]. Other studies have shown estrogen stimulate naïve T cell differentiation to anti-inflammatory Tregs [3]. Following Pivonka et al. [16], we use normalized, weighted Hill repressor and activator functions to capture the inhibitory and stimulatory effects of estrogen in compartment *j*:

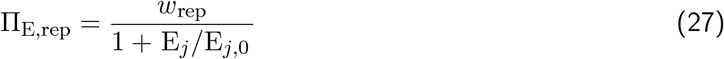

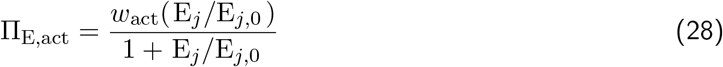

where E_*j*,0_ is the estrogen level corresponding to homeostasis (i.e., the initial/control condition), E_j_ is the estrogen level at time *t*, and *w* are the weight of repression and activation effects.

Adjusting Equations 23–26 to account for estrogen inhibition and stimulation of Th17 and Treg cells:

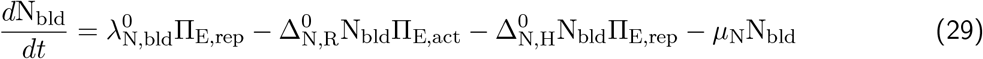

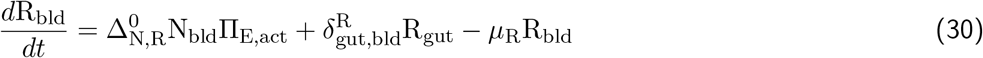

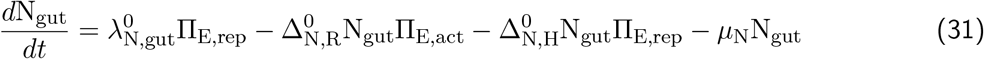

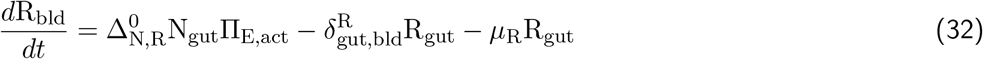

where 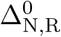 and 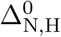 are the differentiation rates of naïve T cells under homeostatic conditions to Treg and Th17 cells. The differentiation rate is modified by changes in estrogen levels E from homeostasis. A decrease in estrogen, as expected during menopause, increases naïve T cell differentiation to proinflammatory Th17 cells (Π_E,rep_ increases) and decreases their differentiation to anti-inflammatory Tregs (Π_E,act_ decreases).

Th17 populations can be tracked similarly to Tregs. Including the effects of estrogen on Th17 cells,

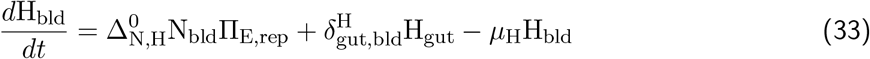

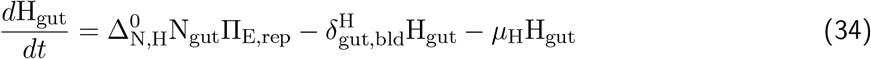

where *µ*_H_ is the degradation rate of Th17 cells and 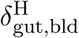 is the migration rate of Th17 cells from the gut to the blood.

### 2.4 Bidirectional T cell migration

To improve the physiological accuracy of the model, we add migration of Tregs and Th17 cells from the blood to the gut. The current model only considers T cell migration from the gut to blood (e.g.,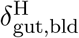). Bidirectional migration can be incorporated with minimal new parameters by adding a coefficient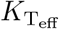, where 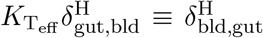 is the migration of Th17 cells from the blood to gut. We can further reduce the number of unknown parameters by defining the coefficient as

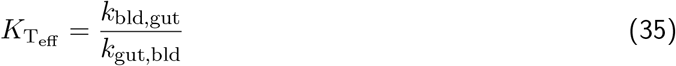

where *k*_*j,i*_ are rates of activated lymphocyte migration from compartment *j* to *i* based on experimental data in the literature (see Section 2.8.4).

We drop the compartment labels on the migration rates by assuming migration rates (*δ*^cell^) are from gut to blood, such that 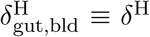 and 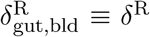 and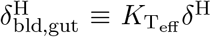. The updated model equations are given in Equations 36–41.

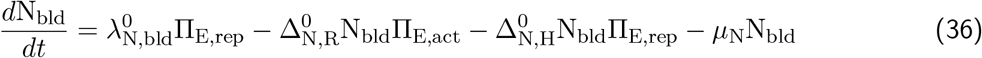

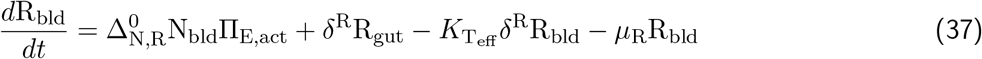

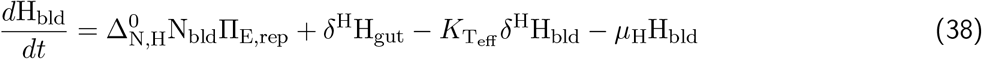

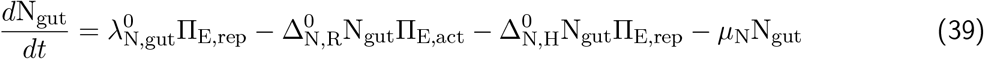

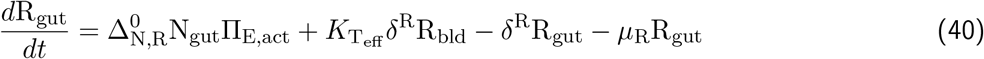

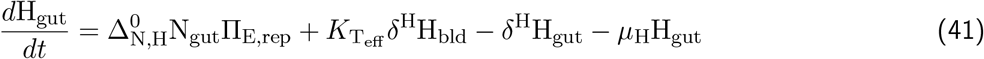

### 2.5 Cell counts from flow cytometry data

Tregs and Th17 levels in the literature are reported as percentages and measured by flow cytometry methods (see Section 2.8.7). Islam et al. [6] converted Treg percentages, or cell fractions, to cell counts by assuming 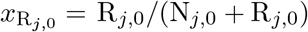. We modify this approach to account for the addition of Th17 cells, such that

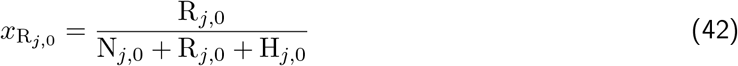

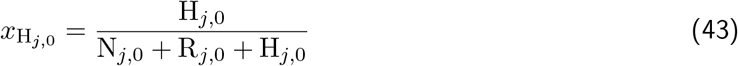

where we assume N_*j*,0_ = 1. To calculate the population of Tregs and Th17 cells from known *x*_R_*j*,0 and *x*_H_*j*,0 values, we rearrange Equations 42 and 43 below.

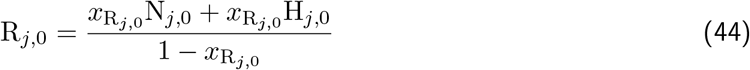

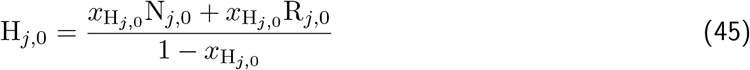

Equations 44 and 45 must be solved simultaneously to calculate R_*j*,0_ and H

### 2.6 Basis variations between compartments

Islam et al. [6] assumed an equal number of naïve T cells in the blood and gut (i.e., N_*j*,0_ = 1). However, the distribution of T cells in the gut and blood is known to vary across the body [12]. Since our model accounts for changes in T cell counts rather than concentration, we can account for the difference in T cell distributions across compartments by assuming different homeostatic basis values for naïve T cells in the blood and gut such that N_bld,0_ ≠ N_gut,0_. Using literature data to calculate the ratio of naïve T cells in the blood and gut 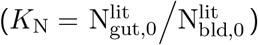 we can set N_bld,0_ = 1 and N_gut,0_ = K_N_N_bld,0_.

### 2.7 Initial conditions from *in vivo* experiments

Initial conditions correspond to the initial values of the homeostatic state, that is the non-treatment condition (here the “sham” condition) where 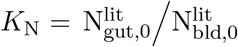 and N_*j*,0_ values are obtained from the literature. N_bld,0_ is then set to 1 and the other initial conditions are obtained from experimental data.

### 2.8 Literature for model parameterization

#### 2.8.1 T cell degradation rates used in modeling literature

T cell degradation rates are commonly set based on existing literature. A narrow look into mathematical models containing T cell dynamics revealed large disparities in the degradation rates of naïve CD4^+^T cells, Treg cells, and Th17 cells (Figure 2). Naïve T cell degradation rates varied 3 orders of magnitude, from 3 × 10^−4^ to 3.3 × 10^−1^ day^−1^. Treg and Th17 degradation rates similarly varied greatly, ranging from 0.001 to 0.5 day^−1^ and 0.007 to 0.5 day^−1^. A closer look reveals that many of these degradation rates are based on experimental data for different cell types or a general assumption from an earlier mathematical model. We can narrow down plausible T cell degradation rates by only considering those based on half-life data from matched experimental data; for example, naïve T cell degradation rates based on experiments measuring CD4^+^naïve T cells. Focusing on the source of these rates, we obtain one mouse and one human study each for naïve T cells and Tregs, and one from mice for Th17 cells. We therefore consider only the experiments from mice to coincide with the aims of this model. Note that both Treg and Th17 studies discussed below were used to obtain degradation rates in Moise and Friedman [17], a rheumatoid arthritis model.

**Figure 2:**
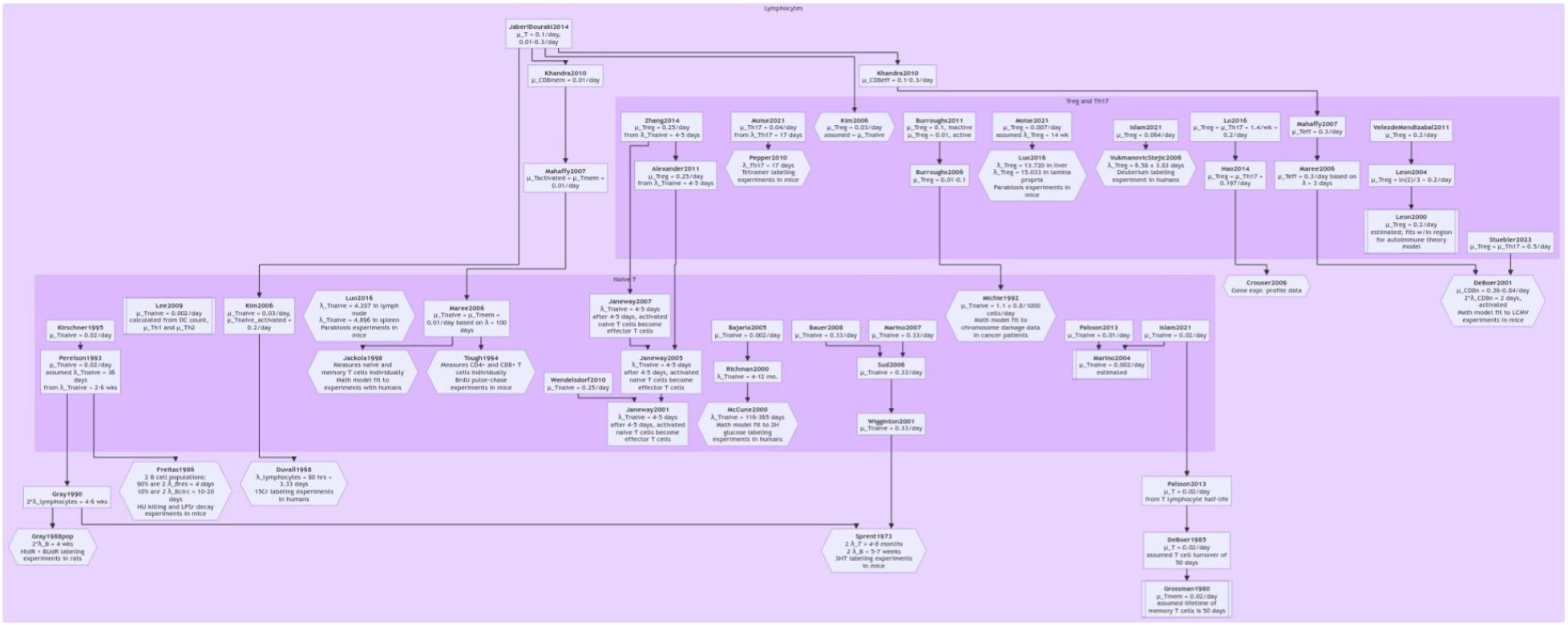
A flowchart of T cell degradation rates used in mathematical models tracing back to their source. The variables *µ*_*i*_, *λ*_*i*_, and 2*λ*_*i*_ represent the degradation rate, half-life, and lifespan of cell type *i*.

#### 2.8.2 Treg degradation rate from experimental data

One of these studies is Luo et al. [14], which includes measurements for both naïve CD4^+^T cell and Treg turnover. Luo et al. [14] employs parabiosis experiments in genetically identical (except at one gene locus; i.e., congenic) C56BL/6 mice are surgically joined to share a circulatory system, then separated after 4 weeks. Cell turnover is then measured in the non-host and half-lives reported. Specifically, total Treg (CD4^+^Foxp3^+^) turnover after disconnection is measured for the liver and lamina propria, while naïve T cell (CD4^+^Foxp3^−^CD62L^hi^CD44^lo^), resting Treg (CD4^+^Foxp3^+^CD62L^hi^CD44^lo^) and activated Treg (CD4^+^Foxp3^+^CD62L^lo^CD44^hi^) turnovers are reported for the spleen and lymph nodes. The half-life of total Tregs is 13.72 − 15.03 days and this range encompasses the half-lives of activated Tregs, while the half-life of naïve T cells is 4.21 − 4.90 days. These correspond to degradation rates of *µ*_N_ = 0.14 − 0.16*/*day and *µ*_R_ = 0.046 − 0.050*/*day based on Equation 46. The similarity in rates between compartments support for our assumption of identical degradation rates in the gut and blood.

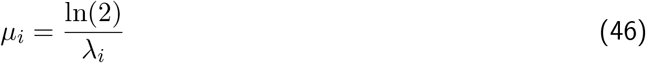

#### 2.8.3 Th17 degradation rate from experimental data

Another study by Pepper et al. [15] measures Th17 cell turnover. Pepper et al. [15] investigates whether CD4^+^memory T cells with antigenic peptides on their MHC class II (pMHCII) molecules exist *in vivo* as T cell subsets such as Th17, Th1, or Th2. The data for Th17 turnover comes from a study investigating the survival of IL-17-producing T cells after infection to determine if they differentiated into memory cells. In this experiment, C56BL/6 mice underwent strain-engineered intranasal infection. After the initial infection response cleared (24 days), the mice were challenged again with infection at various time. Specific pMHCII levels (2W1S:I-A^b^) in the cells were then measured in *ex vivo* stained cells to capture the turnover of IL-17^+^CD4^+^T cells. Since IL-17 cells did not persist after the initial infection response (<0.5% of pMHCII cells were IL-17 producing at day 110), we assume the data for this experiment (Pepper et al. [15] Fig. 4b) is appropriate to use for estimating effector Th17 cell degradation rates.

Moise and Friedman [17] estimates the half-life from the experimental data in Pepper et al. [15] is 17 days. To validate this estimate, we used three different approaches using data extracted with PlotDigitizer. First, we visually estimated that 50% of the initial T cell count occurred at 10–15 days. We also used Equation 47 to calculate the half-life as 14–15 days. The third method used curve-fitting to estimate half-life; using Excel, we fit a power law model to the data and obtained a half-life of 9–12 days. Our results thus indicate the half-life ranges from 9 to 15 days, which at its maximum is 2 days less than that estimated by Moise and Friedman [17]. Using Equation 46, these half-lives corresponding degradation rates of *µ*_H_ = 0.046 − 0.077/day for *λ*_*T h*17_ = 9 − 15 days and *µ*_H_ = 0.041*/*day for *λ*_*T h*17_ = 17 days.

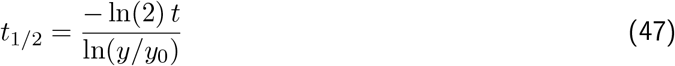

#### 2.8.4 T cell migration parameters from lymphocyte kinetics literature

The migration of lymphocytes has been studied experimentally since the 1950s, evolving from lymphocyte collection to radioactive labeling experiments [11]. However, mathematical modeling of these experiments is necessary to quantify lymphocyte residence times and migration rates.

Our model considers migration of Th17 and Treg cells between the gut and blood compartments. We opt to fit the gut-to-blood migration rates (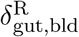 and 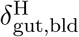) to scaled data from Azam et al. [8], and therefore scale the blood-to-gut rates based on migration rates using using Equation 35. We assume the relative rate of migration is the same for Th17 and Treg cells, both types of effector T cells (i.e., T_eff_).

Literature values for effector T cell migration rates are obtained from the mathematical model of Ganusov and Tomura [11], which are fitted to experimental data from Sprent [18]. In the experimental study, thoracic duct lymphocytes (TDL) from male CBA mice were labeled with ^125^IUdR, injected into male CBA × C56BL/6 mice, and labeled cells measured after sacrifice at various time points. The measured ^125^IUdR cells correspond to activated effector T cells. Of note, the model assumed migration involving the spleen and gut compartments (MLN and intestines) are gamma-distributed, and so they are modeled with *k* sub-compartments. Moreover, the estimated MLN residence times are poorly constrained due to the low number (0.6-2.9%) of cells migrating into the compartment over the experiment duration. With these limitations in mind, we assume the migration rates between gut and blood are a sum of those reported for the MLN and intestines as indicated in Table 4. Standard deviations are approximated based on *n* = 3 mice used in the ^125^IUdR experiments [18].

**Table 4:** Parameter estimates from Ganusov and Tomura [11] model fitted to Sprent [18] data of activated TDL migration in mice. Data are mean ± SD, where SD is calculated using the 95% confidence interval (bracketed values) and *n* = 3.

| Compartment ($j$) | $\delta_{\text{bld},j} (\text{h}^{-1})$ | $\delta_{j,\text{bld}} (\text{h}^{-1})$ |
| --- | --- | --- |
| MLN | $0.7 \pm 0.6$ (0.3-1.6) | $0.1 \pm 0.4$ (0.6-2.9) |
| Intestine | $5.5 \pm 1.9$ (2.9-7.3) | $0.1 \pm 0$ (7.2-12.8) |
| Gut = MLN + Intestine | $6.2 \pm 2.0$ | $0.2 \pm 0.4$ |

From Equation 35, the rate of T cell migration from the blood to gut is given by 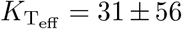 times greater than the rate of migration from the gut to blood.

#### 2.8.5 *In vivo* OVX experiments investigating T cell changes

For our model, we looked to the literature to source experimental data for the change in Th17 and Treg levels in the gut and blood due to OVX. Only four studies include both Th17 and Treg levels in both the gut (PP or MLN) and blood (SP) compartments: Azam et al. [8], Gu et al. [19], Sapra et al. [20], Piao et al. [21]. Except for Piao et al. [21], which used female rats, these studies were conducted on female mice. Gu et al. [19] used C57BL/6J mice, while Azam et al. [8], Sapra et al. [20] both used BALB/c mice.

#### 2.8.6 Systematic search for experimental T cell data

In our search, we sought original research articles studying how OVX in female rodents affected gut inflammation and T cell levels in the gut (PP, MLN, intestines), blood (spleen, serum), and/or bone (Figure 3). Three databases were searched: PubMed/MEDLINE, Embase, and Web of Science. The search query used in PubMed leverages the approach described in Hooijmans et al. [22] to find all available animal experiment studies.

**Figure 3:**
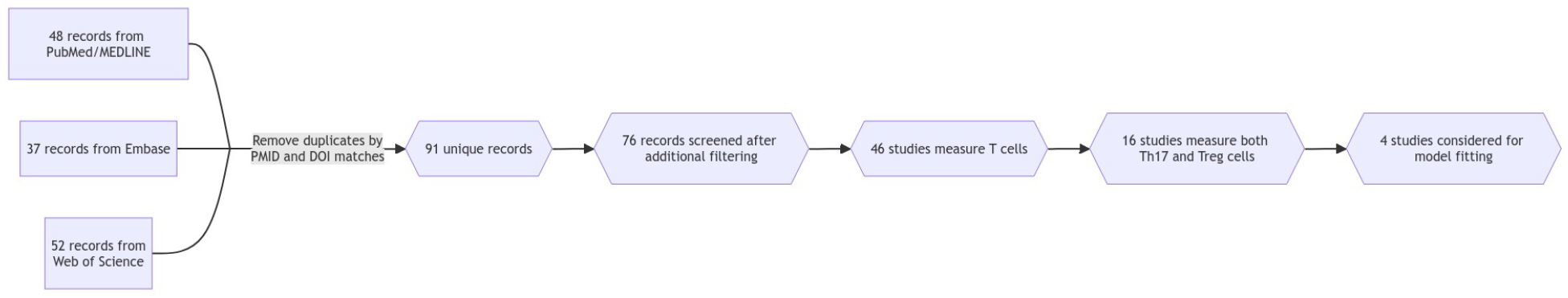
A flowchart summarizing the process for identifying *in vivo* experiments with T cell measurements in the gut and blood in control and OVX rodents. From the unique database records, we further filtered down to remove 3 review articles, 2 articles unavailable in English, 2 *in vitro* studies, and 1 primate study. The remaining 76 records were screened for relevance and excluded if the studies did not investigate gut-immune-bone axis or measure variables of interest. Another 18 articles were excluded for lack of Th17 or Treg measurements, followed by another 10 with only Tregs and 2 with only Th17. Given our model contains 2 compartments, the gut and blood, we would ideally find studies with Tregs and Th17 cell measurements in both compartments. Only 6 of the 16 studies that measured both Th17 and Treg cells included measurements in 2 compartments and only 4 of these 6 studies measured T cell levels in the gut and blood.

**Figure 4:**
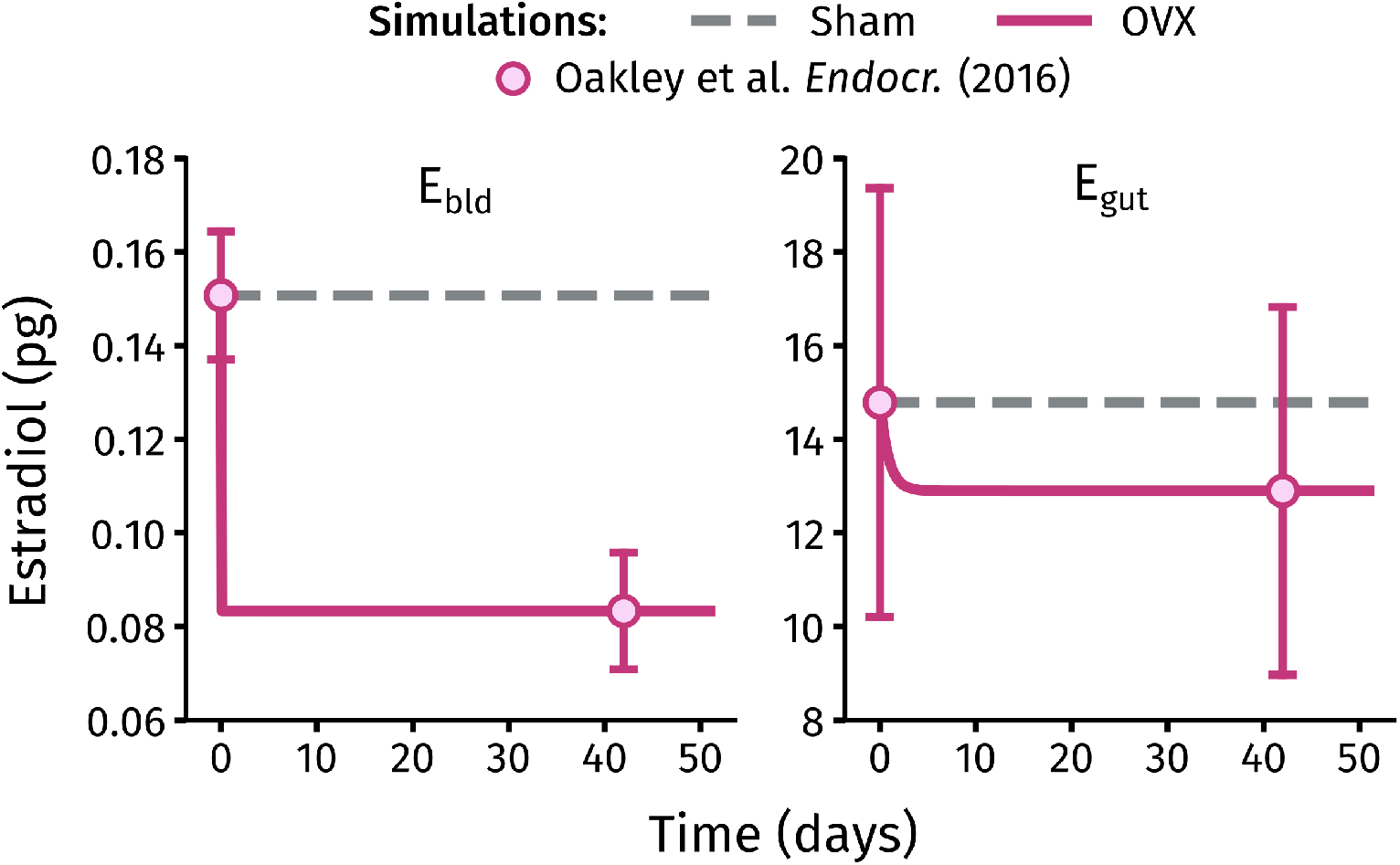
Total estrogen levels in the blood and gut over time under sham and OVX surgery conditions.

#### 2.8.7 Flow cytometry methodology and interpretation

Flow cytometry results are most accurately interpreted when all sample preparation, antibodies/staining, and gating strategies are listed in detail. Sample preparation details can explain why certain gates (e.g., live/dead gating) may be skipped, and help contextualize interpretation of cytokine and transcription factor measurements. The gating strategy and analysis description is also necessary to understand the reported cell percentages. While publications often provide the final dot plot and corresponding percentages, these values typically match final analysis results (e.g., results in bar graphs and table) even when the axis labels of the dot plot and final analysis results seem to contradict each other. Results should explicitly indicate what “percent of” they are reporting, e.g. CD4^+^IL-17^+^cells as a percent of CD4^+^cells or CD4^+^IL-17^+^/CD4^+^, rather than as vague percentages (e.g., Th17 (%)). Without detailed gating descriptions or percentage descriptors, the raw flow cytometry files are needed for re-analysis to allow for accurate interpretation of results.

Gu et al. [19] provides an insufficient description of their flow cytometry methodology that reduces the interpretability of their results. No gating strategy is included in Gu et al. [19], so we must make assumptions to understand and interpret their results. We first assume a standard FSC-SSC gate is used to eliminate debris from the cell count. Since Gu et al. [19] does not mention any viability dyes or markers, we assume no such gate is included. We use the provided list of markers and fluorochromes to assume Th17 cells were gated as follows: (1) SSC/FSC, (2) CD4^+^/FSC, and finally (3) IL-17^+^/CD4^+^. So we assume Tregs were gated by SSC/FSC, CD4^+^/FSC, and then CD25^+^Foxp3^+^/CD4^+^. Together this would indicate the reported Th17 and Treg values are the percentage of IL-17^+^and CD25^+^Foxp3^+^cells out of measured CD4^+^cells. The lack of viability testing or gating indicates the reported T cell values are overestimated due to the presence of dead cells. There is also no mention of techniques necessary for the accurate detection of cytokines and transcription factors; that is, there is no mention of protein transport inhibitors, buffers, nor viability/functionality stimulation.

Azam et al. [8] outlines their antibodies, sample preparation protocol, and flow cytometry gating strategy. Single-cell solutions are prepared, followed by cytokine stimulation (ionomycin + PMA), treatment with protein transport inhibitors, and finally antibody staining. The gating strategy for Tregs and Th17 cells both begin with SSC/FSC, then SSC/CD4^+^, followed by Ror*γ*t^+^/CD4^+^for Th17 cells and Foxp3^+^/CD4^+^for Tregs. So, Th17 and Treg cells are reported as the percentage of Ror*γ*t^+^and Foxp3^+^cells out of measured CD4^+^cells. No mention of viability dyes or testing indicates the reported T cell values are overestimated due to the presence of dead cells.

Sapra et al. [20] outlines their antibodies, sample preparation protocol, and flow cytometry gating strategy. Samples were incubated with anti-CD3 and anti-CD28 monoclonal antibodies, followed by *in vivo* differentiation of Tregs – stimulated with anti-IL-4, anti-IFN-*γ*, TGF-*β* 1 and IL-2 – and Th17 cells – stimulated with anti-IL-4, anti-IFN-*γ*, TGF-*β*-1, IL-6, and IL-23. After 4 days, cells were harvested and stained. The gating strategy is reported to follow Dar et al. [23], which consists of first gating for cells (SSC/FSC), then CD4^+^cells, followed by Foxp3^+^/CD4^+^and Ror*γ*t^+^/CD4^+^for Treg and Th17 cells. So, Th17 and Treg cells are reported as the percentage of Ror*γ*t^+^and Foxp3^+^cells out of measured CD4^+^cells. No mention of viability dyes or testing indicates the reported T cell values are overestimated due to the presence of dead cells.

Piao et al. [21] outlines their antibodies and sample preparation protocol, but not their flow cytometry gating strategy. The lack of a gating strategy requires assumptions to interpret their results. For sample preparation, separate samples of 2 × 10^7^ cells were prepared for Treg and Th17 cell analysis. Samples for Treg detection were treated with transcription factor buffer and then stained (anti-CD4 and anti-Foxp3). Samples for Th17 detection were just stained with anti-CD4 and reportedly anti-CD17 antibodies; however, given the lack of PE-conjugated CD17 antibodies available from eBioscience, we assume an error in the publication such that an anti-IL-17 antibody was used rather than anti-CD17. As with data from Gu et al. [19], we assume a standard FSC-SSC gate is used to eliminate debris from the cell count. However, since no dyes are mentioned, we assume no viability sorting preceded or followed the FSC-SSC gating. We then assume samples are gated for CD4^+^cells, followed by IL-17^+^and Foxp3^+^cells, such that Th17 and Treg cells are reported as the percentage of IL-17^+^and Foxp3^+^cells out of measured CD4^+^cells. We note the discrepancy of this assumption against the y-axis of Figure 2, which would imply the Th17 and Treg cells are reported as the percentage of IL-17^+^CD4^+^and Foxp3^+^CD4^+^cells out of splenocytes and MLN. The number of splenocytes and MLN cells would then be 2 × 10^7^ cells (initial sample size) assuming no SSC/FSC gate, or fewer if there was a SSC/FSC gate.

#### 2.8.8 T cell counts from flow cytometry fraction data

The primary dataset we use for model calibration is from Azam et al. [8] due to the availability and clarity in their Treg and Th17 fractional data in the PP, MLN, and SP.

#### 2.8.9 *In vivo* data of serum and gut estrogen levels

A systematic search of the literature for *in vivo* estrogen data was attempted but proved less effective than other strategies, including ad hoc searches and citation-relation exploration via ResearchRabbit. Our searches returned few relevant papers [7, 24, 25]. Khaw et al. [24] measured sham and OVX female mouse estradiol levels in various lymph nodes, but not gut-associated lymph nodes. More promising was Park et al. [25], which measured estradiol levels in the serum, ovaries, MLN, PP, and ileum. However, these measurements were in female piglets and therefore non-ideal for our mouse-based model. We ultimately found an earlier study from Dr. Ko’s lab at the University of Illinois at Urbana-Champaign, Oakley et al. [7], that met our criteria: *in vivo* experiments in female mice with estradiol measurements under sham and OVX conditions in the serum and gut-related tissues.

The estradiol levels in Oakley et al. [7] are measured by ELISA and reported as pg/mg for tissue and pg/ml for serum. Estradiol levels are measured in the gonads, serum, spleen, ileum, MLN, and PP of female and male mice. The concentration of estradiol is highest in female gonads (Oakley et al. [7] Fig 3B) as one might instinctively expect. Estradiol concentrations are lower in the serum than gut-associated tissues. However, total estradiol content was higher in female gut-associated tissues than ovaries. Comparing healthy and OVX mice (Oakley et al. [7] Fig 5E), there is a significant decrease in serum estradiol concentration for OVX mice but no significant difference in gut tissue estradiol concentration. Oakley et al. [7] therefore concludes gut estradiol levels are not driven by ovarian secretion of estrogen and instead driven by factors within the intestinal environment.

**Figure 5:**
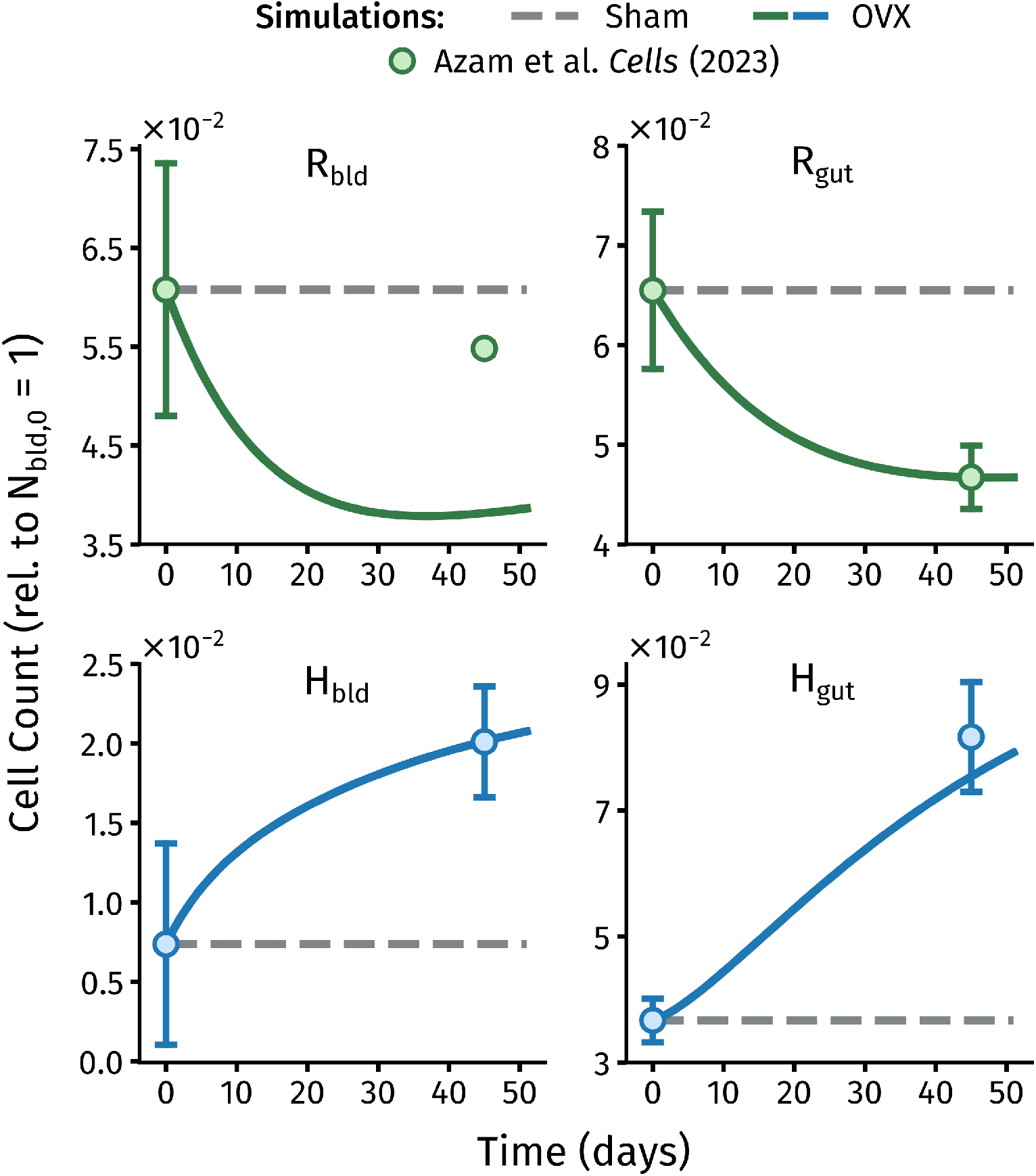
Normalized T cell levels in the blood and gut over time under sham and OVX surgery conditions.

#### 2.8.10 Gut compartment mass and blood compartment volume

Since Oakley et al. [7] only reports total estrogen levels for healthy mice and not OVX mice, we need the blood compartment volume and gut compartment masses to calculate total estrogen levels in OVX mice. We calculate the MLN and PP compartment masses using

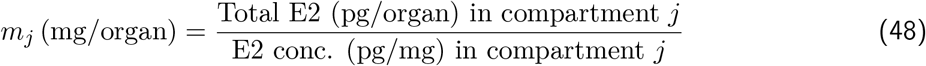

where the total estrogen and estrogen concentrations (conc.) are extracted from Oakley et al. [7] Fig 5B-C with PlotDigitizer.

The total blood compartment volume is calculated from alternate literature sources using Equation 49. A general blood volume for mice is 76-80 ml/kg [26]. For mouse weight, we use the mean weight of 78 *±* 2 g from Azam et al. [8], as this will approximate the estrogen levels associated with the T cell data.

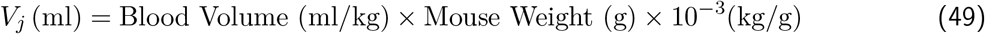

#### 2.8.11 Total estrogen content from concentration data

Our model tracks total estrogen levels to match with our T cell model, which tracks cell counts rather than concentrations. Oakley et al. [7] reports total estrogen levels for healthy but not OVX mice. Using the volume and masses calculated above, we calculate total estrogen levels per compartment for sham and OVX mice using

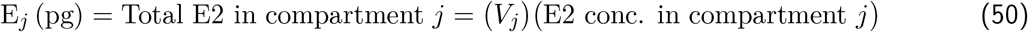

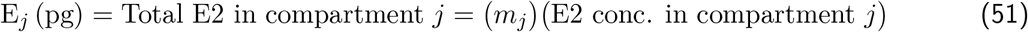

where Equation 50 applies to serum estrogen levels and Equation 51 applies to tissue estrogen levels. Furthermore, we validate the above approach by comparing the calculated total estrogen levels in WT mice to the measured levels from Oakley et al. [7]. Table 5 shows that the total estrogen contents are lower (1.69 and 13.1) than the measured values (1.82 and 16) and the error is higher. However, the calculated levels are within a standard deviation of the measured estrogen content, so this approach is reasonable.

**Table 5:** Comparison of calculated and measured total estrogen in WT mice. Data are mean *±* SD.

| Compartment | $m_j$ (mg) | Measured $E_j$ (pg) | Calculated $E_j$ (pg) |
| --- | --- | --- | --- |
| PP | $3.58 \pm 0.68$ | $1.82 \pm 0.20$ | $1.69 \pm 0.36$ |
| MLN | $24.38 \pm 5.32$ | $16 \pm 1.89$ | $13.10 \pm 4.56$ |

To calculate the gut compartment estrogen content, we sum the MLN and PP compartments. Note that ileum data is excluded due to the lack of ileum T cell measurements in Azam et al. [8], which only includes T cell fractions in the MLN and PP. Due to data availability, the blood compartment estrogen content is calculated from serum estrogen concentration, while blood compartment T cell levels are from the spleen. The data and intermediate calculations are given in Table 6 and the final gut and blood compartment estrogen levels are given in Table 7.

**Table 6:** Calculated total estrogen values from concentration data in Oakley et al. [7]. Data are mean *±* SD. ^*^Serum compartment reported as pg/ml and *V*_*j*_ (ml).

| Condition | Compartment | Conc. E (pg/mg)* | $m_j$ (mg)* | Calculated $E_j$ (pg) |
| --- | --- | --- | --- | --- |
| WT | PP | $0.47 \pm 0.05$ | $3.58 \pm 0.68$ | $1.69 \pm 0.36$ |
| WT | MLN | $0.53 \pm 0.15$ | $24.38 \pm 5.32$ | $13.10 \pm 4.56$ |
| WT | Serum | $0.08 \pm \text{NA}$ | $1.79 \pm 0.16$ | $0.15 \pm 0.01$ |
| OVX | PP | $0.61 \pm 0.20$ | $3.58 \pm 0.68$ | $2.2 \pm 0.81$ |
| OVX | MLN | $0.44 \pm 0.12$ | $24.38 \pm 5.32$ | $10.73 \pm 3.84$ |
| OVX | Serum | $0.05 \pm 0.006$ | $1.79 \pm 0.16$ | $0.08 \pm 0.01$ |

**Table 7:** Total estrogen content in the gut and blood compartment. These are the estrogen data points we use to calibrate the estrogen model. Data are mean *±* SD.

| Condition | Compartment | Calculated $E_j$ (pg) |
| --- | --- | --- |
| WT | Gut | $14.79 \pm 4.58$ |
| WT | Blood | $0.151 \pm 0.014$ |
| OVX | Gut | $12.90 \pm 3.93$ |
| OVX | Blood | $0.083 \pm 0.012$ |

## 3 Results and Discussion

Results for simulations of sham and OVX conditions are shown in Figures 4 and 5.

## 4 Abbreviations and Symbols

**Abbreviations**

Foxp3: forkhead box protein 3 14, 15
IFN-*γ*: interferon gamma 15
IL-17: interleukin-17 12, 14, 15
IL-2: interleukin-2 15
IL-23: interleukin-23 15
IL-4: interleukin-4 15
IL-6: interleukin-6 15
MLN: mesenteric lymph node 13, 15–17
OVX: ovariectomy 3–5, 13–19
PP: Peyer’s Patch 13, 15–17
Ror*γ*t: RAR-related orphan receptor gamma two 15
SP: spleen 13, 15
TGF-*β*: transforming growth factor beta 15
Th17: T helper 17 4–6, 8–15
Treg: regulatory T cell 4–15
WT: wild type 16, 17

## State Variables

| Symbol | Description | Unit | Page List |
| --- | --- | --- | --- |
| B | butyrate | $\mu\text{M}$ | 6, 7 |
| E | estrogen | ng | 3–5, 8–11, 16, 17 |
| H | T helper 17 cell | count | 6, 8–13 |
| N | naïve T cell | count | 6–12, 20 |
| R | regulatory T cell | count | 6–13 |

## Parameters

| Symbol | Description | Unit | Page List |
| --- | --- | --- | --- |
| $\Delta$ | differentiation rate | $\text{day}^{-1}$ | 6–10 |
| $\delta$ | migration rate | $\text{day}^{-1}$ | 4, 5, 7–10, 13 |
| $\gamma$ | *in vitro* to *in vivo* factor | - | 6, 7 |
| $\Pi$ | Hill function | - | 9, 10 |
| $K_N$ | relative amount of naïve T cells in the blood vs gut | - | 8, 10 |
| $K_{T_{\text{eff}}}$ | relative migration of T-cells from the blood to gut | - | 8–10, 13 |
| $\lambda$ | formation or production rate | cells/day | 4–10 |
| $\mu$ | elimination or apoptosis rate | $\text{day}^{-1}$ | 4, 5, 7–10, 12 |
| $\tau$ | state of ovarian function | - | 4, 5 |
| $w$ | weight parameter | - | 8, 9 |

## Subscripts and Superscripts

| Symbol | Description | Page List |
| --- | --- | --- |
| 0 | initial | 4–11 |
| $j$ | compartment | 4–10, 16, 17 |
| bld | blood | 4–11, 13 |
| gut | gut | 4–11, 13 |
| rep | repression or inhibition | 8–10 |
| act | activation or stimulation | 8–10 |
| cond | treatment condition reflecting state of ovarian function | 4, 5 |
| lit | value from the literature | 10 |
| ovx | ovariectomy (OVX) surgery condition | 4, 5, 8 |
| sham | sham surgery condition | 4, 5, 8 |

## Acknowledgments

This work was supported by National Institutes of Health grants R35 GM133763, R21 AG077640, and R15 AT010725 and the University at Buffalo. AML was also partially supported by the Arthur A. Schomburg Fellowship Program of the University at Buffalo and the NSF Graduate Research Fellowship Program.

## References

[1] Khosla, S., Oursler, M. J., and Monroe, D. G. Estrogen and the skeleton. Trends Endocrinol. Metab., 23(11):576–581, 2012.

[2] Tai, P., Wang, J., Jin, H., Song, X., Yan, J., Kang, Y., Zhao, L., An, X., Du, X., Chen, X., Wang, S., Xia, G., and Wang, B. Induction of regulatory T cells by physiological level estrogen. J. Cell. Physiol., 214(2):456–464, 2007.

[3] Li, J.-Y., Chassaing, B., Tyagi, A. M., Vaccaro, C., Luo, T., Adams, J., Darby, T. M., Weitzmann, M. N., Mulle, J. G., Gewirtz, A. T., Jones, R. M., and Pacifici, R. Sex steroid deficiency–associated bone loss is microbiota dependent and prevented by probiotics. J. Clin. Investig., 126(6):2049–2063, 2016.

[4] Smith, B. J., Hatter, B., Washburn, K., Graef-Downard, J., Ojo, B. A., El-Rassi, G. D., Cichewicz, R. H., Payton, M. E., and Lucas, E. A. Dried plum’s polyphenolic compounds and carbohydrates contribute to its osteoprotective effects and exhibit prebiotic activity in estrogen deficient C57BL/6 mice. Nutrients, 14(9):1685, 2022.

[5] Artoni de Carvalho, J. A., Magalhães, L. R., Polastri, L. M., Batista, I. E. T., de Castro Bremer, S., Caetano, H. R. d. S., Rufino, M. N., and Bremer-Neto, H. Prebiotics improve osteoporosis indicators in a preclinical model: systematic review with meta-analysis. Nutr. Rev., 81(8):891–903, 2022.

[6] Islam, M. A., Cook, C. V., Smith, B. J., and Ford Versypt, A. N. Mathematical modeling of the gut–bone axis and implications of butyrate treatment on osteoimmunology. Ind. Eng. Chem. Res., 60(49):17814–17825, 2021.

[7] Oakley, O. R., Kim, K. J., Lin, P.-C., Barakat, R., Cacioppo, J. A., Li, Z., Whitaker, A., Chung, K. C., Mei, W., and Ko, C. Estradiol synthesis in gut-associated lymphoid tissue: leukocyte regulation by a sexually monomorphic system. Endocrinology, 157(12):4579–4587, 2016.

[8] Azam, Z., Sapra, L., Baghel, K., Sinha, N., Gupta, R. K., Soni, V., Saini, C., Mishra, P. K., and Srivastava, R. K. Cissus quadrangularis (Hadjod) inhibits RANKL-induced osteoclastogenesis and augments bone health in an estrogen-deficient preclinical model of osteoporosis via modulating the host osteoimmune system. Cells, 12(2):216, 2023.

[9] Stanczyk, F. Z., Archer, D. F., and Bhavnani, B. R. Ethinyl estradiol and 17β-estradiol in combined oral contraceptives: pharmacokinetics, pharmacodynamics and risk assessment. Contraception, 87 (6):706–727, 2013.

[10] Yoshinaga, K., Hawkins, R. A., and Stocker, J. F. Estrogen secretion by the rat ovary in vivo during the estrous cycle and pregnancy. Endocrinology, 85(1):103–112, 1969.

[11] Ganusov, V. V. and Tomura, M. Experimental and mathematical approaches to quantify recirculation kinetics of lymphocytes, pages 151–169. Springer International Publishing, Cham, 2021.

[12] Chen, Z., Huang, A., Sun, J., Jiang, T., Qin, F. X.-F., and Wu, A. Inference of immune cell composition on the expression profiles of mouse tissue. Sci. Rep., 7(1):40508, 2017.

[13] den Braber, Ineke and Mugwagwa, T., Vrisekoop, N., Westera, L., Mögling, R., Bregje de Boer, A., Willems, N., Schrijver, E. R., Spierenburg, G., Gaiser, K., Mul, E., Otto, S., Ruiter, A. C., Ackermans, M., Miedema, F., Borghans, J. M., de Boer, R., and Tesselaar, K. Maintenance of peripheral naive T cells is sustained by thymus output in mice but not humans. Immunity, 36(2): 288–297, 2012.

[14] Luo, C. T., Liao, W., Dadi, S., Toure, A., and Li, M. O. Graded Foxo1 activity in Treg cells differentiates tumour immunity from spontaneous autoimmunity. Nature, 529(7587):532–536, 2016.

[15] Pepper, M., Linehan, J. L., Pagán, A. J., Zell, T., Dileepan, T., Cleary, P. P., and Jenkins, M. K. Different routes of bacterial infection induce long-lived TH1 memory cells and short-lived TH17 cells. Nat. Immunol., 11(1):83–89, 2010.

[16] Pivonka, P., Zimak, J., Smith, D. W., Gardiner, B. S., Dunstan, C. R., Sims, N. A., Martin, T. J., and Mundy, G. R. Model structure and control of bone remodeling: a theoretical study. Bone, 43 (2):249–263, 2008.

[17] Moise, N. and Friedman, A. Rheumatoid arthritis - a mathematical model. J. Theor. Biol., 461: 17–33, 2019.

[18] Sprent, J. Fate of H2-activated T lymphocytes in syngeneic hosts. I. Fate in lymphoid tissues and intestines traced with 3h-thymidine, 125i-deoxyuridine and 51chromium. Cell. Immunol., 21(2): 278–302, 1976.

[19] Gu, J., Zheng, Y., Yang, H., Li, Y., Liu, S., Wu, Y., Ren, L., Yu, Y., and Long, Y. Cistanche deserticola polysaccharide regulated the gut microbiota-SCFAs-Th17/Treg cell axis and ameliorated the inflammation of postmenopausal osteoporosis. J. Functional Foods, 109:105811, 2023.

[20] Sapra, L., Dar, H. Y., Bhardwaj, A., Pandey, A., Kumari, S., Azam, Z., Upmanyu, V., Anwar, A., Shukla, P., Mishra, P. K., Saini, C., Verma, B., and Srivastava, R. K. Lactobacillus rhamnosus attenuates bone loss and maintains bone health by skewing Treg-Th17 cell balance in Ovx mice. Sci. Rep., 11(1):1807, 2021.

[21] Piao, J., Park, J. S., Hwang, D. Y., Son, Y., and Hong, H. S. Substance P blocks ovariectomy-induced bone loss by modulating inflammation and potentiating stem cell function. Aging, 12(20): 20753–20777, 2020.

[22] Hooijmans, C. R., Tillema, A., Leenaars, M., and Ritskes-Hoitinga, M. Enhancing search efficiency by means of a search filter for finding all studies on animal experimentation in PubMed. Lab Anim., 44(3):170–175, 2010.

[23] Dar, H. Y., Shukla, P., Mishra, P. K., Anupam, R., Mondal, R. K., Tomar, G. B., Sharma, V., and Srivastava, R. K. Lactobacillus acidophilus inhibits bone loss and increases bone heterogeneity in osteoporotic mice via modulating Treg-Th17 cell balance. Bone Rep., 8:46–56, 2018.

[24] Khaw, Y. M., Anwar, S., Zhou, J., Kawano, T., Lin, P.-C., Otero, A., Barakat, R., Drnevich, J., Takahashi, T., Ko, C. J., and Inoue, M. Estrogen receptor alpha signaling in dendritic cells modulates autoimmune disease phenotype in mice. EMBO Rep., 24(3):e54228, 2023.

[25] Park, C. J., Kim, H., Jin, J., Barakat, R., Lin, P.-C., Choi, J. M., and Ko, C. J. Porcine intestinal lymphoid tissues synthesize estradiol. J. Vet. Sci., 19(4):477–482, 2018.

[26] van Zutphen, L. F. M., Baumans, V., and Beynen, A. C., editors. Principles of laboratory animal science, volume 26. Elsevier, Portland: Copyright Clearance Center., rev. ed. edition, 2002.

